# Assessment of Generative *De Novo* Peptide Design Methods for G Protein-Coupled Receptors

**DOI:** 10.64898/2026.02.26.708415

**Authors:** Hannes Junker, Clara T. Schoeder

## Abstract

G protein-coupled receptors (GPCRs) play an ubiquitous role in the transduction of extracellular stimuli into intracellular responses and therefore represent a major target for the development of novel peptide-based therapeutics. In fact, approximately 30% of all non-sensory GPCRs are peptide-targeted, representing a blueprint for the design of *de novo* peptides, both as pharmacological tools and therapeutics. The recent advances of deep learning-based protein structure generation and structure prediction offer a multitude of peptide design stategies for GPCRs, yet confidence metrics rarely correlate with experimental success. In the context of peptides, this problem is exacerbated due to the lack of elaborate tertiary structures in peptides, raising the question of whether this is due to inadequate sampling or insufficient scoring. In this two-part benchmark, we addressed this question by first simulating the validation process of 124 unique known GPCR-peptide complexes using AlphaFold2 Initial Guess, Boltz-2 and RosettaFold3. We then assessed the peptide sampling capabilities of the respective generative methods BindCraft, BoltzGen and RFdiffusion3. Our results indicate that current design pipelines primarily suffer from significant confidence overestimation for misplaced peptides in the validation phase across all three prediction methods. We further highlight occurrences of significant memorization in both prediction as well as generation of peptides. While all generative methods sample backbone space sufficiently, their simultaneous sequence generation remains subpar and can be partially recovered through the use of ProteinMPNN. Taken together, our benchmark offers guidance for the design of peptides specifically using deep learning-based pipelines.

**Autor summary:** Deep learning-based protein design is revolutionizing computational biology and development of such tools is progressing rapidly with increasing attention from both academic and non-academic institutions. Their applicability and performance is often assessed from an all-purpose objective, with implicit bias towards larger protein-protein interactions. Due to their size, peptides therefore present an edge case where performance is known to decrease compared to larger, more structured proteins. Here, we present a benchmark specifically for the deep learning-based design of peptides targeting G protein-coupled receptors (GPCRs), a major therapeutic drug target family, assessing the generation of novel GPCR-targeting peptides and the validation of these designs separately. Our results show that generative methods sample potential peptide placements and orientations sufficiently but validation fails to differentiate valid from invalid designs, indicating that the so-called ‘scoring problem’ remains unsolved. Although focusing on a specific use-case, our results are generalizable to the broader field of protein design. Consequently, it can offer guidance for peptide-specific design applications and can contribute to the development and improvement of new methods.

## Introduction

Following the groundbreaking impact of AlphaFold2 (1) the field of deep learning-based computational structural biology is advancing rapidly, enabling a plethora of novel applications in the development of pharmaceuticals (2–6). On the one hand, this includes improved methods for the prediction of biomolecular complexes, e.g. AlphaFold3 (7), Boltz-2 (8),Chai-1 (9) or RosettaFold3 (10). On the other hand, generative models such as RFdiffusion (11), BindCraft (12), RFdiffusion3 (13), Chai-2 (14) or BoltzGen (15) enable the generation of entirely new proteins with atomic precision. In the field of generative protein design, these methods often intertwine in various pipelines where designed proteins are validated through structure prediction. These pipelines may be composed of separate steps for backbone and sequence design with subsequent validation, as in the case with RFdiffusion-based approaches (16), or through inversion of and iterative refinement with structure prediction models for simultaneous sequence and backbone design, as demonstrated with BindCraft. Recently, development shifted towards all-atom generative design resulting in simultaneous generation of sequence and backbone in one inference run as implemented in RFdiffusion3 or BoltzGen. It is, however, still recommended to validate designs through structure prediction with RosettaFold3 or Boltz-2, respectively.

Generally, it is widely understood that current structure prediction methods excel for structured proteins or, in turn, are less confident when predicting unstructured loops. Likewise, generative design models often generate highly structured proteins with high secondary structure content, bundling into rigid tertiary structures (17,18). To counter this, the first version of RFdiffusion, for instance, provides specific weights to reduce helical content, while BindCraft provides settings with softened AlphaFold2 confidence thresholds in order to allow the design of disordered peptides. Similarly, RFdiffusion3 biases generation towards unstructured protein segments via a ‘is_non_loopy’ flag.

Peptides are a steadily growing branch of therapeutics with benefits over small-molecules such as lower toxicity or higher potency and selectivity (19–23). Due to their limited size of 50 or fewer amino acids they encompass many of the unfavorable features that make them difficult to work with in deep-learning based contexts. Peptides are often unable to form elaborate tertiary structures with structural stability mainly driven by shorter α-helical stretches or the formation of β-hairpins that can unravel once binding to a target protein is initiated (24). Endogenously, peptides act as hormones, growth-factors or neurotransmitters and play a substantial role in G protein-coupled receptor (GPCR) signaling, with approximately 30% of non-sensory GPCRs being targeted by peptide endogenously (25). GPCRs are the largest group of proteins in the human genome and are major pharmacological targets with approximately 34% of FDA-approved drugs targeting GPCRs (25,26). Structurally, GPCRs are composed of a highly conserved seven transmembrane helical bundle whose orthosteric binding site is formed by the extracellular pocket formed by these helices. The pocket can further be occluded by extracellular loops surrounding it. Upon binding of a ligand an allosteric cascade facilitates the opening of the intracellular G protein binding site (27–29). From an *in silico* design perspective, GPCRs are a complicated target as the orthosteric pocket is often too tight to allow the formation of secondary structures. At the same time, endogenous peptides are highly affine and selective for their cognate receptor where minor changes in sequence composition, charge or backbone orientation can eliminate the allosteric communication between extracellular ligand binding site and intracellular G protein binding site (30–32). For *de novo* peptides to mimic or compete with endogenous peptides, similar specificity is mandatory.

As high-throughput production and testing of *de novo* peptides is significantly less feasible compared to *de novo* proteins robust filters and selection criteria are imperative. While prediction accuracies specifically of the entire ligand-GPCR-G protein complex have been assessed previously for AlphaFold2 and AlphaFold3 (33,34), a *de novo* design-centric benchmark for GPCRs is missing. Taking into account the inherent limitations of current prediction and generation tools with respect to short, unstructured polypeptides we understand this to be a question of sampling vs. scoring. With a limited pocket volume to work with, do generative design models sample the space of possible peptides adequately and propose plausible peptide candidates? Subsequently, are prediction methods capable of assigning these putative peptides an appropriate confidence score to differentiate a probable candidate from an improbable candidate? We sought to address these questions in a GPCR-specific two-part benchmark that reflects typical *de novo* design pipeline setups. With regard to scoring, we investigated how accurate and confident AlphaFold2 Initial Guess (AF2IG) (35), Boltz-2 and RosettaFold3 (RF3) predict the placement of endogenous peptides relative to their cognate receptor for 124 unique receptor-peptide complexes. We chose these three prediction methods as they support templating, where a structure template guides the prediction of a target receptor while the respective network is tasked with the placement of the peptide. Secondly, for three representative peptide-receptor complexes, we generated 10000 putative peptides with BindCraft, BoltzGen and RFdiffusion3 each with the goal of mimicking the native peptide. Through structural deviation to the reference peptide we examine the explorational power of each generative design model. Here, we selected BindCraft, BoltzGen and RFdiffusion3 since these can generate both backbone and sequence and are therefore independent of another inverse-folding model such as ProteinMPNN (36). Furthermore, this allowed for each generative method to be evaluated both for prediction and generation, thus being cross-validated orthogonally.

## Results

### Collection of GPCR-peptide complexes

In order to investigate whether AF2IG, Boltz-2 and RF3 are able to predict the binding mode of GPCR-peptide complexes, we collected a set of crystal structures by filtering the GPCRdb (37) for ‘peptide’ or ‘protein’ ligands, amounting to a total of 414 complexes. Although the latter is technically outside the scope of peptides, their binding mechanism to GPCRs is often similar to peptide-GPCR interactions in that they also have a peptide-like disordered terminus entering the orthosteric binding pocket. From here on both peptide and protein ligands were collectively referred to as peptides. The dataset was further refined to only contian non-redundant GPCR-peptide dimers whose peptides are gapless and otherwise do not contain any non-canonical amino acids or functional groups (see Methods for details regarding processing and filtering). This results in a dataset of 124 unique GPCR-peptide dimers (see Supplementary Table S1). The final dataset consists of 76 peptides binding to 58 class A receptors and 31 peptides binding to 14 class B1 receptors, reflecting the promiscuous many-to-many binding relationship that is characteristic for many GPCRs and their endogenous peptides (Fig. 1A). The majority of peptides are between five and 21 amino acids long and bound to class A receptors, with the shortest consisting of only three amino acids (pseudoallergen receptor MRGPRX2 in complex with three resolved residues of substance P; PDB ID *7vdm* (38)). Peptides of length 27-61 were mostly hormones binding to class B1 receptors. Peptides of length 62-73 were mostly chemokines or chemotactic cytokines. Eight receptors binding to nanobodies or single domain antibodies were not excluded from the dataset as their binding is mediated by loops and can be regarded as the intersection of peptides and protein binders (Fig. 1B).

**Fig 1:**
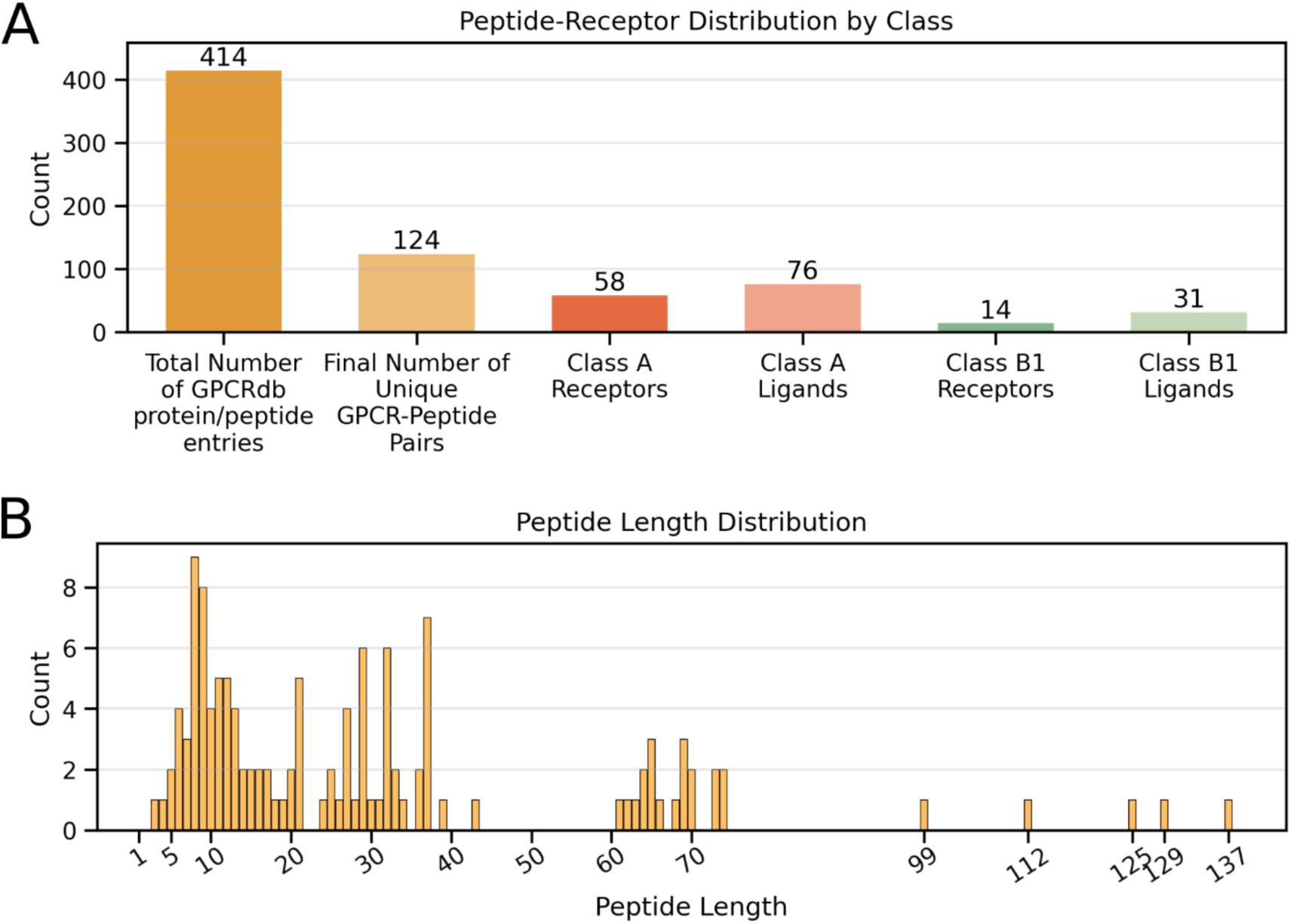
Dataset distribution. **A** Receptor and peptide distribution split by GPCR class. A receptor can occur multiple times with different peptides and a peptide can bind to multiple receptors. **B** Distribution of peptide lengths. Peptides of length 25-61 bound to receptors of class B1; Peptides of length 65-74 mostly consist of peptides binding to class A Chemokine and Chemotactic receptors; lengths >74 consisted of nanobodies and single domain antibodies.

### Benchmarking prediction of GPCR-peptide complexes

The first aim of this benchmark was to determine the capability of AF2IG, Boltz-2 and RF3 to reproduce known GPCR-peptide binding modes. To this end, the crystal structure of each of the 124 receptors was provided as a template to Boltz-2 and RF3. In case of AF2IG, the dimer of the receptor and its peptide was the input, where the coordinates of the receptor are used to initialize the prediction. For the prediction of the peptides multiple sequence alignments (MSAs) were not provided for either of the prediction methods with the rationale to mimic *de novo* peptide prediction. Each receptor-peptide dimer was consequently predicted 50 times using a different seed for each run.

To measure the structural deviation of predictions to crystal structure reference, the predicted dimer was aligned with the crystal structure by superimposing the receptors. Next, the DockQ score (39) was calculated for the predicted peptide to the reference peptide. Analogous to the categorization of the DockQ authors, we will refer to predictions as ‘incorrect’ (DockQ < 0.23), ‘acceptable quality’ (0.23 ≤ DockQ < 0.49), ‘medium quality’ (0.49 ≤ DockQ < 0.8) and ‘high quality’ (DockQ ≥ 0.80).

### Peptide predictions were seed-dependent

All three prediction methods were, to some extent, capable of reproducing some of the receptor-peptide dimers in the dataset, however, with significantly different success rates. Collectively, over all 50 predictions for all 124 dimers, AF2IG achieved a median DockQ score of 0.03, with 70.65% of all predictions being ‘incorrect’ (DockQ < 0.23) and only 7.15% ‘high quality’ predictions (DockQ > 0.8) (Fig. 2A). This is marked by a median iRMSD of 12.2 Å and 68% of predictions with fnat ≤ 0.1 (Fig. 2B and C). Both RF3 and Boltz-2 outperformed AF2IG in this regard. RF3 achieved a median DockQ score of 0.41 and exhibited a slightly bimodal DockQ score distribution with 36.31% of predictions being classified as ‘incorrect’, 24.22% high quality’ predictions and the remaining 39.46% of predictions falling in the range 0.23 < DockQ < 0.8, i.e. ‘acceptable quality’ and ‘medium quality’ predictions. Boltz-2 achieves the best median DockQ score of 0.56 with 62.16% of predictions being of ‘medium quality’ or better. Although the majority of Boltz-2 predictions showed a low iRMSD (median of 2.2 Å), a small number of peptide predictions were drastically misplaced with an iRMSD of 66.26 Å.

**Fig. 2:**
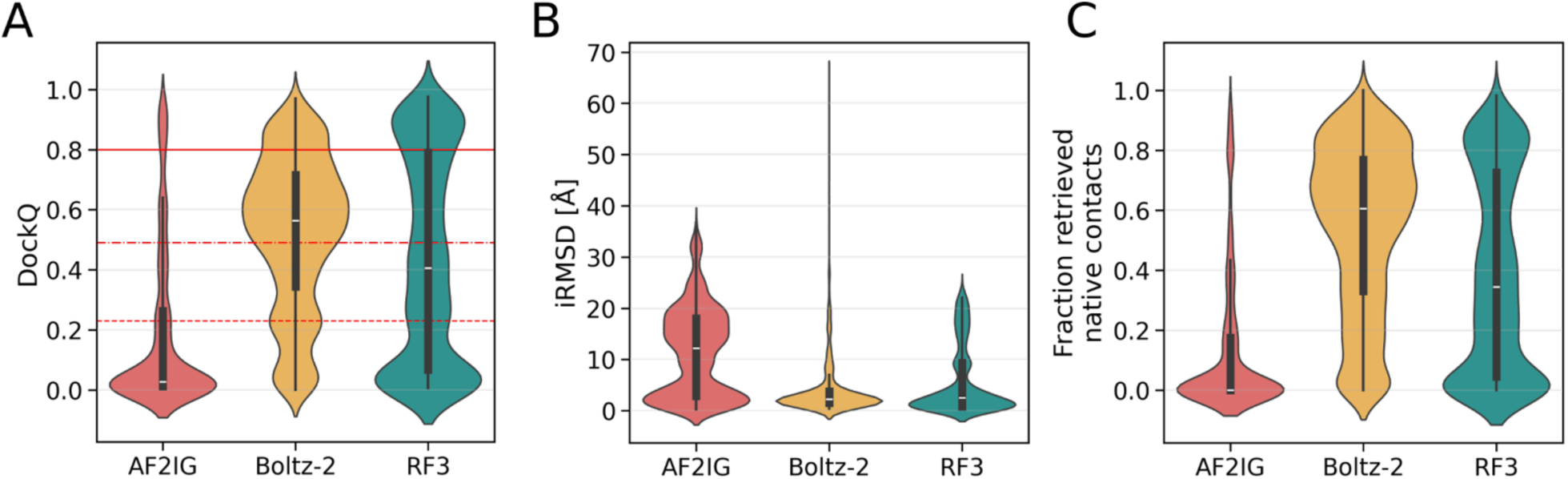
Collective structural deviation of 50 predictions per prediction method of all 124 receptor-peptide complexes. **A** DockQ score distributions. A DockQ score < 0.23 (dashed red line) is considered “Incorrect”, a score 0.23 ≤ DockQ < 0.49 (dash-dotted red line) is considered “Acceptable”, a score 0.49 ≤ DockQ < 0.80 (solid line) is considered “Medium” and DockQ ≥ 0.80 is considered as “High quality”. **B** Distributions of RMSD of interface residues (iRMSD). **C** Distributions of fractions of retrieved native contacts (fnat).

Inspecting the 50 predictions per receptor-peptide dimer revealed that using different seeds seemed to result in vastly different prediction qualities (Fig. 3). AF2IG was least affected by such seed inconsistencies with some exceptions, e.g. PDB IDs *8flu* (40) or *6b3j* (41) where, depending on the seed, the prediction of the peptide ligand can either be ‘incorrect’, with a DockQ score as low as 0.03 or ‘high quality’ with a DockQ score as high as 0.85. For some peptides, Boltz-2 and RF3 produced similarly wide-ranging DockQ scores across different seeds and both methods were generally prone to seed-dependent DockQ score spread. In this regard, Boltz-2 produced varying DockQ scores for most of the peptides which often resulted in Boltz-2 outperforming AF2IG and RF3 when comparing the highest scoring seeds per method. Generally, a variance in DockQ score seemed to be attributed to a similar variance in the fraction of retrieved native contacts (Fig. 2) and less to variance in iRMSD or LRMSD, which was unsurprising regarding the fact that a relatively minor peptide backbone shift can result in the disruption of most contacts. Additionally, the peptide length influences DockQ scores slightly for AF2IG and RF3, while it has no impact on DockQ scores for Boltz-2 (Supplementary Fig. S2). Interestingly, the three prediction methods performed differently from peptide to peptide. For example, for PDB ID *7w53* (42), RF3 predicted the peptide with little structural deviation to the crystal reference (median DockQ = 0.81). Conversely, Boltz-2 and AF2IG predictions of that peptide only achieved a median DockQ score of 0.24 and 0.12, respectively. For PDB ID *8f7w* (43), a peptide of similar length (8 AA compared to 7 AA in *7w53*), Boltz-2 achieved a median DockQ score of 0.83, while AF2IG and RF3 achieved a median DockQ score of only 0.24 and 0.43, respectively.

**Fig. 3:**
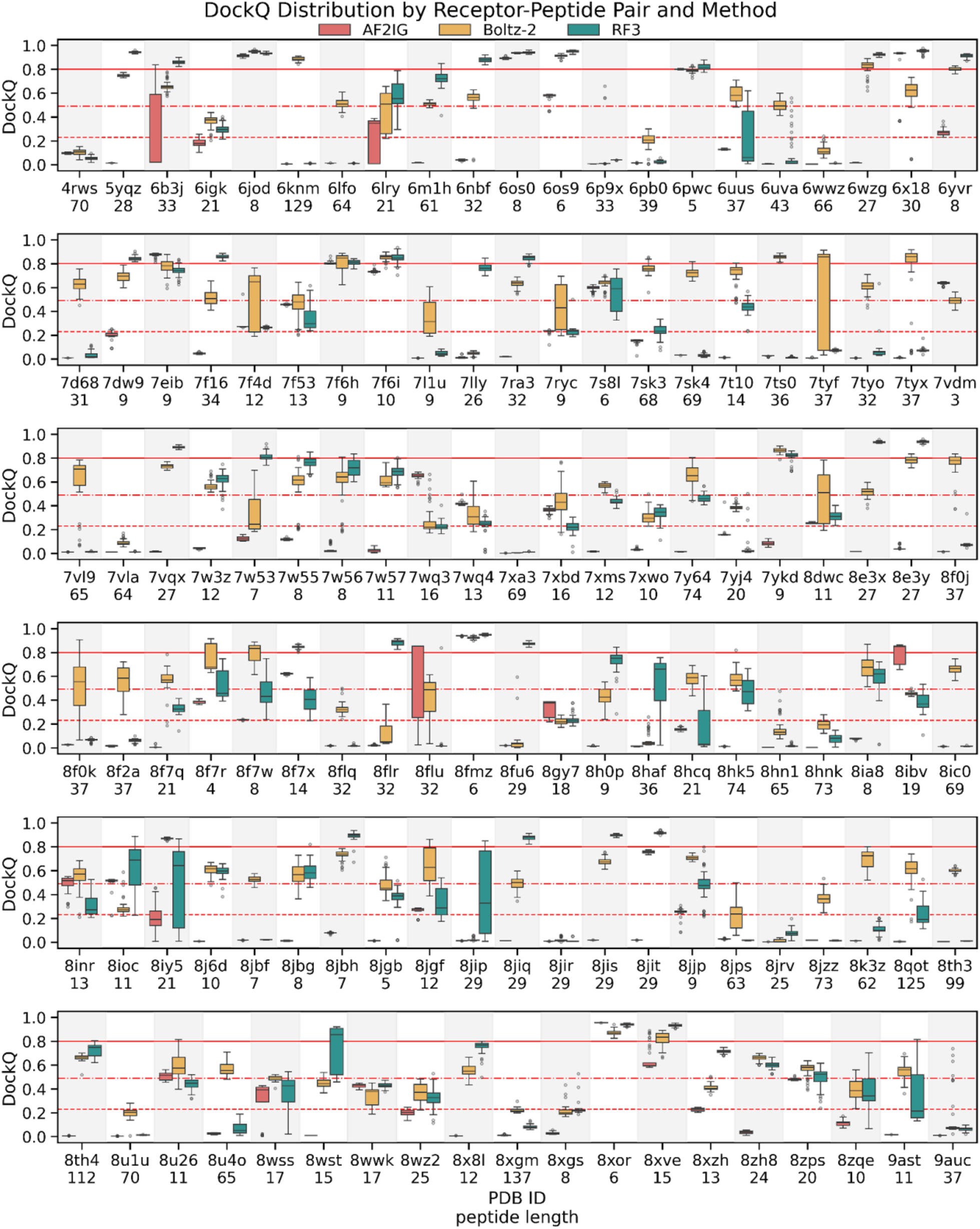
DockQ score distribution over 50 predictions for each of the 124 receptor-peptide pairs and for each prediction method (blue: AF2IG, orange: Boltz2, green: RF3). A DockQ score < 0.23 (dashed red line) is considered “Incorrect”, a score 0.23 ≤ DockQ < 0.49 (dash-dotted red line) is considered “Acceptable”, a score 0.49 ≤ DockQ < 0.80 (solid red line) is considered “Medium” and DockQ ≥ 0.80 is considered as “High quality”. Numbers below PDB IDs refer to the length of the respective peptide.

### Prediction confidence scores weakly correlated with structural deviation

As a typical *de novo* peptide or protein design procedure usually involves filtering based on different confidence metrics, we investigated whether the three prediction methods could distinguish between correctly and incorrectly predicted peptides. We opted to focus primarily on confidence scores derived from different subsets of the predicted aligned error matrix (PAE) as other confidence metrics such as the predicted template modeling score (pTM) or the newer predicted docking error (PDE) could not be extracted or calculated in a sufficiently comparable manner. More specifically, we selected the peptide intra-chain PAE subset (termed “PAE peptide” from here on), the peptide-receptor inter-chain PAE subset (also known as PAE-interaction but termed “inter-chain PAE” from here on to avoid ambiguity) and the ipSAE score (44) as most suitable to describe confidences for different parts of the peptide-receptor complex (Fig. 4). Here, we find that inter-chain PAE for predictions performed with AF2IG (and to a lesser extent with RF3) was most suitable to distinguish correct from incorrect peptide predictions with Spearman’s correlation of −0.829 and −0.894, respectively, between confidence and actual structural deviation. Furthermore, the average pLDDT for the peptide chain shows slight correlation with DockQ score for Boltz-2 and RF3 predictions but not for AF2IG predictions (Supplementary Fig. S3). An overall trend with the other PAE confidences is over-estimation, where incorrect predictions exhibit the same PAE or ipSAE score as correct predictions, signifying a large number of false positives. This is most prominent with Boltz-2, where medium to low quality predictions were indistinguishable from high quality predictions through confidence alone. Conversely, there appears to be a ceiling effect where predictions with high DockQ score were rarely assigned a bad PAE/ipSAE score, in other words a low number of false-negatives. Specifically selecting entries from the PAE matrix that correspond to interchain pairs with ≤ 10 Å distance is entirely unsuitable to differentiate accurate from inaccurate placements (Supplementary Fig. S3).

**Fig. 4:**
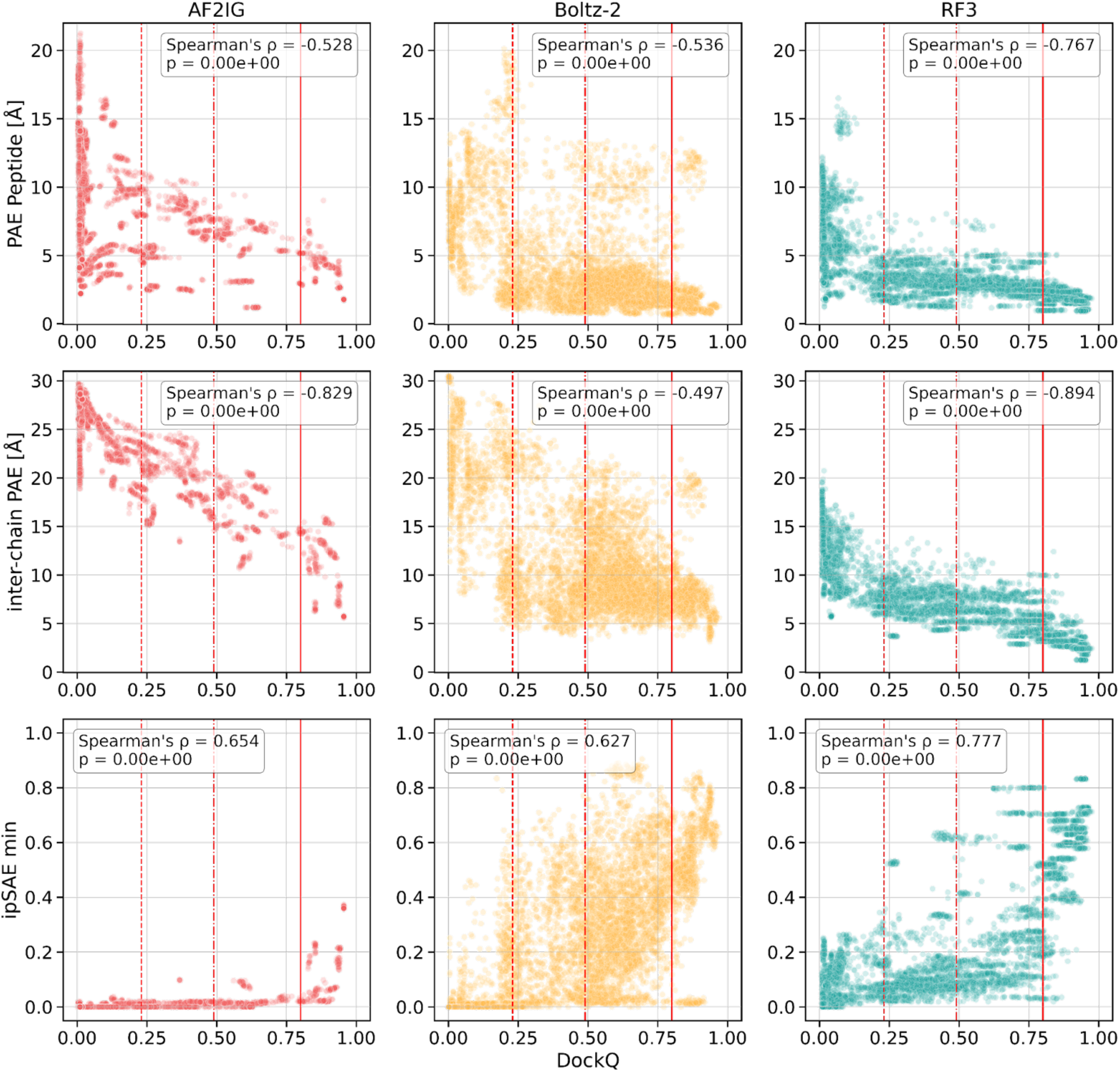
PAE based confidences compared to DockQ scores by method for all predictions. Top row: Averaged PAE values for the peptide chain. Middle row: PAE inter-chain, calculated as the average of all interchain PAE values. Bottom row: iPSAE min calculated as in (44). A DockQ score < 0.23 (dashed red line) is considered “Incorrect”, a score 0.23 ≤ DockQ < 0.49 (dash-dotted red line) is considered “Acceptable”, a score 0.49 ≤ DockQ < 0.80 (solid red line) is considered “Medium” and DockQ ≥ 0.80 is considered as “High quality”

### Prediction diversity illustrated by Endothelin receptor complexes with different peptides

The Endothelin receptor type B (ET_B_) was present in the dataset a total of five times with different peptides and reflects the many-to-many relationship that is characteristic for GPCR-peptide interactions. For the purpose of this benchmark, ET_B_ was especially interesting as it binds to multiple peptides that are virtually identical in structure but differ in sequence composition (Fig. 5A). This resembles a typical scenario in *de novo* design where a number of sequences generated with an inverse-folding method, e.g. ProteinMPNN, would be predicted and selected based on confidence metrics. Concretely, the Endothelin-binding peptides investigated here were the endogenous agonists endothelin-1 (PDB ID *8iy5* (45)) and endothelin-3 (PDB ID *6igk* (46)), as well as sarafotoxin S6b (PDB ID *6lry* (47)), a snake venom that was evolved to mimic the endogenous peptides. Furthermore, these three peptides were cyclic by formation of two cyclic disulfide bridges which was a desirable condition in peptide design. Lastly, it is worth noting that structures of all three peptide-receptor complexes were included in training of AF2, Boltz-2 and RF3, respectively. As depicted in Fig. 5B, predicting each peptide 50 times with different seeds encompasses many of the characteristics mentioned previously, where we observed inconsistencies when comparing the three methods with each other for each peptide individually as well as when comparing the outputs of each individual method across all peptides with each other. Across all three peptides, AF2IG is unable to reproduce the correct peptide placement with correspondingly high inter-chain PAE scores. In contrast, Boltz-2 reproduced endothelin-1 virtually perfectly with a narrow distribution in both DockQ and inter-chain PAE scores, yet it demonstrated stronger seed-dependent DockQ variance for endothelin-3 and sarafotoxin S6b. The corresponding inter-chain PAE scores, however, only increased slightly compared to the near-perfect predictions of endothelin-1, highlighting that incorrectly placed peptides are not necessarily detectable through confidence values. Lastly, RF3 demonstrated significant seed-dependent spread in DockQ scores for endothelin-1, ranging from incorrect to high-quality predictions. For endothelin-3 it was unable to predict correct peptide placements and for sarafotoxin S6b the structural prediction was predominantly of medium quality. This diversity was strongly contrasted by high prediction confidences with narrow spread across all three peptides, demonstrating that for RF3, incorrect predictions are indistinguishable from correct predictions when considering PAE confidences alone.

**Fig. 5:**
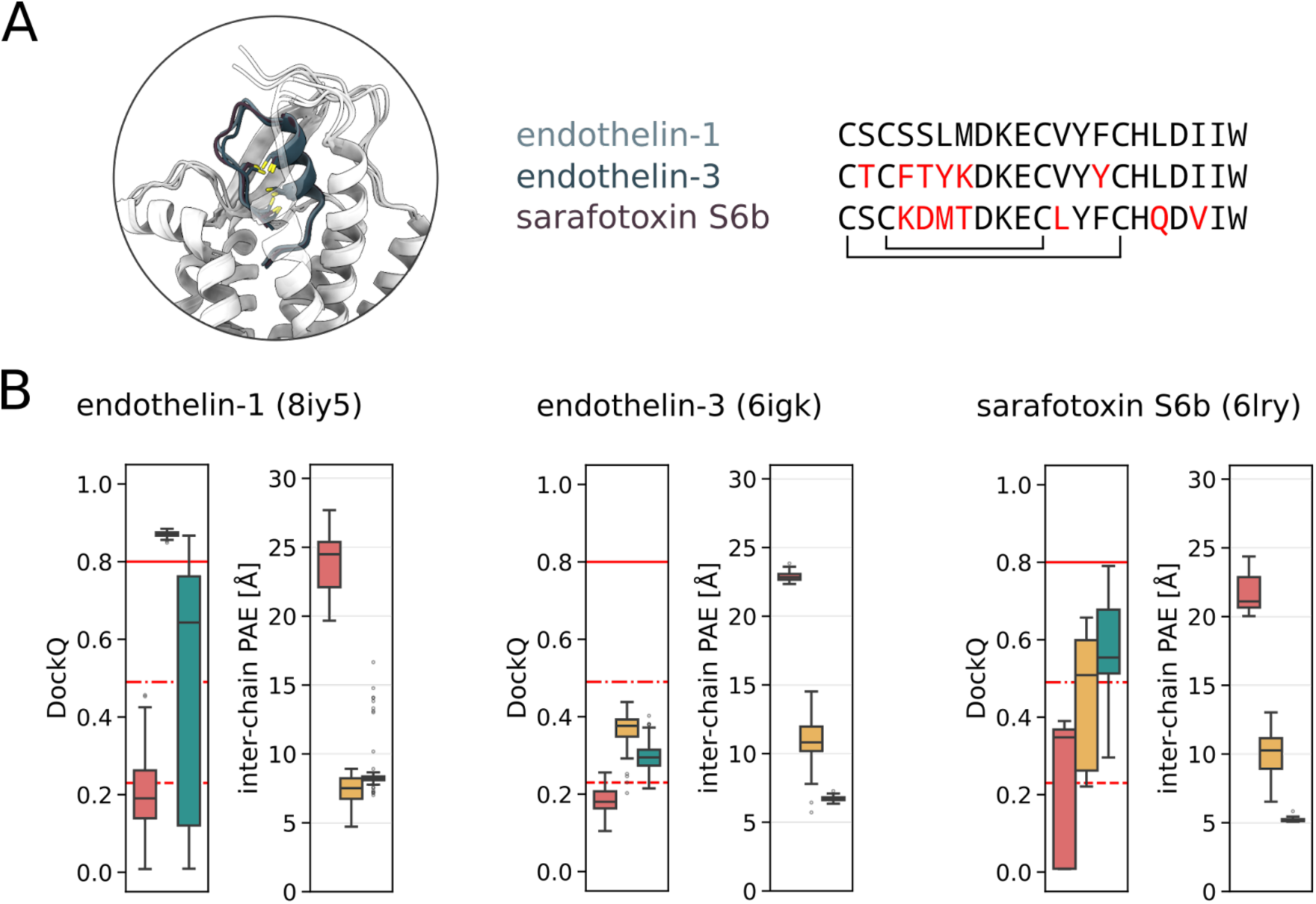
Different Endothelin receptor type B (ET_B_) peptides as example for the discrepancy between DockQ variance and confidence. **A left** Superposition of ET_B_ in complex with endogenous ligands endothelin-1, endothelin-3 and snake toxin sarafotoxin S6b. **A right** sequence alignment of endothelin-1, endothelin-3 and sarafotoxin S6b. Marked in red are residues that differ from endothelin-1. Brackets below the sequences indicate disulfide bridges. **B** DockQ scores and inter-chain PAE values for 50 predictions of endothelin-1, endothelin-3 and sarafotoxin S6b.

### Comparing the Sampling of BindCraft, BoltzGen and RFdiffusion3

Next, we sought to investigate the peptide sampling capabilities of BindCraft, BoltzGen and RFdiffusion3 as these correspond by design to the three structure prediction tools investigated above. Three receptors from our dataset served as target while the complex with its native peptide serves as structural reference: Angiotensin II type 2 (AT_2_) receptor with its endogenous peptide Ang II (PDB ID *6jod* (48), peptide length 8 amino acids), Endothelin receptor type B (ET_B_) in complex with Sarafotoxin S6b (PDB ID *6lry* (47), peptide length 22 amino acids), and Nociceptin (NOP) receptor in complex with its endogenous Nociceptin peptide (PDB ID *8f7x* (43), peptide length 14 amino acids). For each reference complex, three to six residues closest to the pocket-intruding peptide terminus were selected as hotspot residues to guide generation into the orthosteric binding pocket (See Supplementary Fig. S4). In order to mimic the native peptide the *de novo* generation was restricted to exactly the length of the respective reference peptide. With each method 10000 designs were generated per target (see Methods for detailed settings). In the context of GPCRs and their spatially restricted orthosteric binding pocket two criteria were of interest with regard to space sampled through generative design: a) the distance of the designed peptide to the provided hotspots to measure if it reached into the pocket deep enough to address functionally relevant receptor residues and b) Cα-RMSD of the generated peptide to the native peptide to measure the overall structural diversity (Fig. 6). These two distances must be considered together to differentiate peptides placed inside the binding pocket from peptides placed outside the binding pocket. A typical error in GPCR-targeting peptide design is that the designed peptide is placed on the outside of the helix bundle and in contact with membrane-facing residues. Such cases often result in a short hotspot distance, e.g. if a peptide contacts a helix from the outside that holds a hotspot. For our examples such misplaced peptides typically have an RMSD > 20 Å. Peptides that border the opening of the binding pocket but do not enter it are typically marked with a hotspot distance >10 Å in combination with an RMSD of ∼15 Å or lower. Peptides that are ‘flipped’, i.e. enter with the opposite terminus than the reference peptide have a high RMSD but a short hotspot distance.

**Fig 6.**
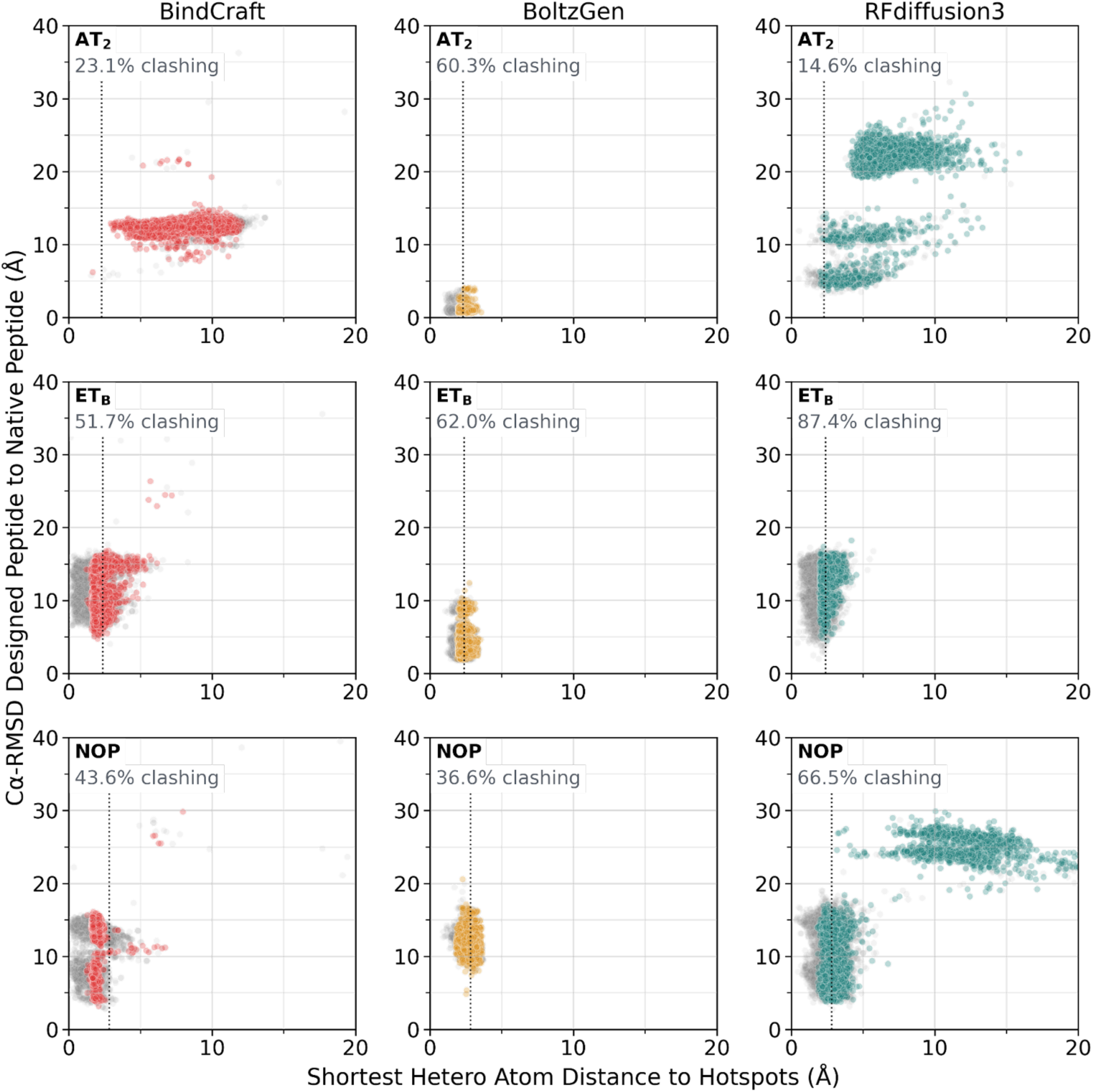
Structural assessment of 10000 generated peptides for each target receptor with each method. Peptide generation was restricted to native peptide length (AT_2_: 8, ET_B_: 22, NOP: 14) and guided by hotspots. Shortest distance to hotspots refers to the closest hetero atom to any hetero atom of the given hotspots. The dotted line signifies the shortest distance of the respective reference peptide to the hotspots. Outliers beyond 40 Å RMSD and 20 Å hotspot distance not shown for visual clarity. Gray dots are peptides with one or more steric clashes. Clashes were determined with the Rosetta PerResidueClashMetric without minimization/relaxation.

Through these two distances we observed that all methods, apart from BindCraft peptides targeting the AT_2_ receptor, were able to place peptides inside the binding pocket. BindCraft and RFdiffusion3 show the broadest diversity in peptide placement inside the binding pocket, effectively sampling the entire binding pocket from shallow entries at the opening of the binding pocket to peptides entering deep into the pocket. However, especially with RFdiffusion3, this came with a large portion of unusable designs that were placed outside the binding pocket and in contact with membrane-spanning residues (18.1% of NOP receptor-targeting and 84.5% of AT_2_ receptor-targeting peptides). Conversely, BoltzGen not only placed every of the 30000 peptides inside the binding pocket, but did so with remarkable similarity to the reference peptides. When targeting ET_B_ receptor, where the reference peptide is the aforementioned cyclic Sarafotoxin S6b, 79.8% of generated peptides have an RMSD ≤ 5 Å to the reference, effectively regenerating cyclic orientations identical or similar to Sarafotoxin S6b or the endogenous endothelin peptides. When targeting AT_2_ receptor, every generated peptide had an RMSD < 5 Å with 97.9% of peptides with an RMSD ≤ 1.5 Å. To further take the precision of the side-chain placement into account, we used the Rosetta PerResidueClashMetric and observe that a significant amount of designed peptides were clashing with the receptor, with up to 87.4% of RFdiffusion3-designed peptides targeting ET_B_ receptor showing at least one steric clash.

Lastly, we selected 90 (10 per method and receptor) sequences of non-clashing peptides inside the binding pocket, randomly sampled uniformly along the RMSD axis and validated them with AF2IG, Boltz-2 and RF3 (50 predictions with different seeds) and used DockQ to assess their placement compared to the initial design (Fig. 7). Overall, we observed that BindCraft designs received the highest scores AF2IG (60.9% ‘high quality’) and BoltzGen designs received the highest scores with both RF3 and Boltz-2 (39.2% and 32.1% ‘high quality’, respectively). Similarly, RFdiffusion3 designs were validated best with RF3, albeit significantly less effectively (12.2% ‘high quality’). In order to dissect backbone design and sequence design we generated one ProteinMPNN sequence for each of the 90 backbones and validated those analogously. Here, we observed a reduction in the DockQ high quality category but at the same time an improvement of previously incorrectly placed peptides, effectively recovering the placement through an optimized sequence. In virtually every generation-validation method combination an enrichment of ‘acceptable’ or ‘medium quality’ DockQ scores after ProteinMPNN sequence design is evident. This effect was strongest for RFdiffusion3 designs, where a single ProteinMPNN sequence improves ‘medium quality’ DockQ scores from 23.1% to 34.2% when validated with AF2IG, from 16.9% to 39.2% when validated with Boltz-2 and from 22.3% to 48.1% when validated with RF3.

**Fig. 7:**
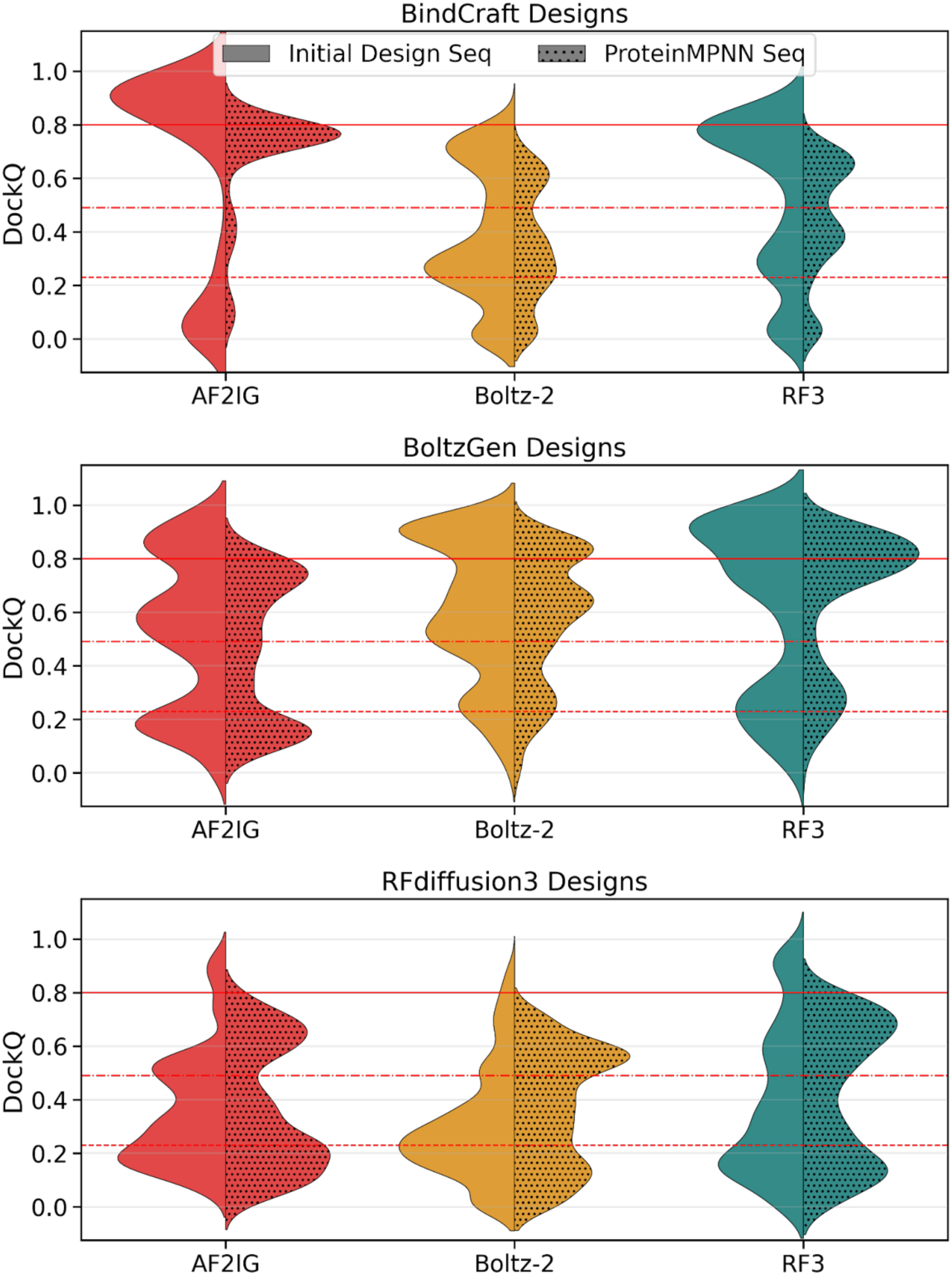
Structural deviations of repredictions of selected designed peptides with AF2IG, Boltz-2 and RF3, respectively. 10 out of 10000 generated peptides per method and reference (90 in total) were selected uniformly along peptide RMSD ≤ 18 Å and with hotspot distance ± 2 Å of reference hotspot distance (see Fig. 6). Clear violins: Sequences from the respective design method. Dotted violins: One randomly selected ProteinMPNN sequence per designed peptide backbone. Repredictions followed the same protocol as previously (50 different seeds, receptor structure given as template, no MSA for the peptide sequence). The respective initially designed peptide served as reference for DockQ in both cases.

## Discussion

Deep learning methods have revolutionized experimental success rates for protein design, however, the transferrability to peptide design for important drug targets has been lacking behind. In the light of frequent release of new tools, re-occuring assessment of their performance on peptide design is imperative. Here, we offer a peptide-GPCR-specific assessment of current possibilities and limitations of recently released methods in structure prediction and generation. The problem of high false positive rates in computational design is not new but due to the lack of references for *de novo* proteins, it is often viewed from the perspective of experimental success (49,50). By simulating a *de novo* peptide design task with known structural references we substantiated the fact that PAE matrix-derived confidence metrics often overestimate and are generally insufficient in differentiating correct peptide placements from incorrect placements. We further highlight that validation of *de novo* peptides is highly seed-dependent, which adds to the large scope of false positive predictions. While these characteristics hold true for all three of the tested prediction methods we showed that each structure prediction method performs differently given specific targets in terms of placement accuracy and associated confidence. This complicates comparability of *de novo* design protocols centered around specific prediction methods as confidence thresholds are not transferable.

While this study focussed on confidence in the form of predicted aligned error, our results align with previously reported inconsistencies with regard to other confidence metrics such as the pTM score for predictions of flexible regions of peptides, which resulted in refined subset scores like actifpTM (51). Generally, it is appreciated that considering confidence score subsets specifically for interacting residues can prove beneficial. In our case, interchain PAE showed a stronger signal than the smaller subset of PAE values used in ipSAE_min and ipSAE_max, while taking only the subset of actually interacting residues, i.e. distance-based PAE, performed worse (Supplementary Fig. S3). This could be attributed to the fact that any confidence metric to some extent reflects coevolutionary information that cannot be solely reduced to interacting residues. At the same time, this might be different for other design targets as GPCRs are structurally and mechanistically highly conserved while the orthosteric binding site requires less conservation to facilitate ligand selectivity.

In this context it is worth noting that no MSAs have been used in our setup. This is a typical trade-off between runtime and accuracy and the number of false positive peptide predictions might decrease if MSAs were included. At the same time, exemplified by three structurally identical peptides targeting the ET_B_ receptor, we identified highly accurate predictions for one specific combination of target and peptide sequence using Boltz-2, which suggests that this combination has been memorized. We want to stress that memorization is likely not unique to Boltz-2. In fact, for all three prediction methods tested here, placement accuracy and confidence significantly decrease for structures from our dataset that were submitted after the respective training cutoffs (Supplementary Fig. 1). Further in-depth analysis is required to investigate the true extent of memorization for any of these methods.

Taking the validation aspect aside, we further demonstrated that all three generative methods were sufficiently capable of sampling the backbone space inside the spatially constrained GPCR binding pocket. Given large enough sampling numbers plausible peptide candidates have been and can be generated. Here, BoltzGen was most precise, however, the results for two of our three examples suggest that this, again, is likely due to memorization. While our results do not show memorization for BindCraft and RFdiffusion3, it cannot be ruled out that these methods can show similar behavior for other targets.

Overall, the number of generations necessary can probably be reduced by fine-tuning the location and number of hotspot residues. Furthermore, we applied each method with near-default settings and each method offers a number of adjustments to balance designability and diversity, which have to be evaluated before designing on a specific target. Generally, we observed that limitations lie primarily in the simultaneous generation of backbone and sequence. Although each method found some suitable sequences for any given backbone, we would like to highlight the strength of ProteinMPNN sequence optimization as a single ProteinMPNN-generated sequence per backbone significantly improved prediction placement of previously incorrect placements. Similarly, in the special case of membrane-bound proteins, a substantial improvement in avoiding placements in the membrane region could be achieved by solubilizing the target protein, either through the use of SolubleMPNN (52) or through rationally mutating the respective positions, e.g. to arginine or lysine.

In all cases, a large portion of the initially designed peptides showed steric clashes, reflecting similar findings for deep learning-based protein-ligand docking methods (53). On that note, it is important to remember that structure prediction and structure generation tools are not equivalent to protein-protein or peptide-protein docking algorithms. While they are extremely powerful in providing a statistical approximation for the placement and orientation of the peptide, they lack the required precision for side-chain interactions. We purposefully excluded sterically clashing designs from further analysis although they might still be plausible candidates if they were subjected to refinement, i.e. in the form of Rosetta minimization or relaxation (54). This ties into the current limitations of confidence-first filters and we strongly suggest incorporating supplemental structural filters for *de novo* peptide design, either through measuring the recovery of hotspot residue contacts, through scaffolding or through comparison with known peptides in combination with physics-informed validations such as the Rosetta interface_ΔG or shape complementarity, similar to BindCrafts filter implementation. In terms of filtering and analysis, BoltzGen also offers a range of metrics to use for filtering, including liability/developability assessments, however, these metrics are based on the unrefined predicted structures. In any case, rigorous post-analysis of *in silico* results prior to experimental testing in order to derive structure-activity relationship hypotheses is time-costly but might be worthwhile.

In conclusion, we demonstrated that BindCraft, BoltzGen and RFdiffusion3 in combination with AlphaFold2 Initial Guess, Boltz-2 and RosettaFold3 remain valuable resources even for hard targets such as peptide design for G protein-coupled receptors. From the current standpoint we suggest rationally motivated peptide generation as opposed to ‘blind’ exploration. Depending on the level of exploration required BoltzGen might be more suitable for narrow sampling while RFdiffusion3 would allow for broader sampling. With its refinement cycles and the inclusion of non-deep learning-based metrics, BindCraft offers a balanced approach between sampling and scoring. To avoid the reliance on a single structure prediction tool for validation, we suggest the use of orthogonal structure validation, e.g. by using both RosettaFold3 and Boltz-2 with a large number of seeds. Overall, by acknowledging their respective weaknesses these tools will be invaluable in the identification of novel GPCR-targeting peptide pharmaceuticals.

## Methods

### Data Availablity Statement

All data relevant to the study are included in the article or uploaded as supplemental information. The receptor dataset as well as the relevant scripts to reproduce structure prediction and generation will be made publicly available upon publication on Zenodo at link https://doi.org/10.5281/zenodo.18755323. Further information and requests for computational and material resources should be directed to and will be fulfilled by the Lead Contact.

### Data collection and curation

We filtered the GPCRdb with ligand type ‘peptide’ or ‘protein’ and downloaded the respective entries from RCSB.org (55). To ensure that any reference peptide was reproducible by all three prediction methods we took following measures: Complexes whose peptides contain gaps or non-canonical amino acids were removed from the dataset. Peptides with unresolved extracellular termini were taken as-is and the sequence used for predictions was truncated accordingly. Any functional groups attached to the binding pocket-intruding terminus, such as NH_2_, were removed. Although these functional groups usually are crucial for GPCR activation, we argue that they play a minor role in the prediction process of the remaining peptide backbone. Additional chains that were neither receptor nor ligand were removed. Finally, to have each receptor-peptide dimer present only once duplicate dimers were filtered for the best crystal structure resolution, resulting in 124 receptor-peptide dimers (Supplementary Table 1).

### Calculation of structural deviation

The DockQ score was chosen over other scores such as RMSD, TM-score (56) or lDDT (57) as they were not fully applicable to any of the dimers in the dataset due to size of the peptides. For instance, since RMSD scales with the number of amino acids, a comparison between peptides of different lengths is hindered. Moreover, the TM-score, although designed to circumvent this problem, is generally applicable to globular proteins. More specifically, due to the design of the scaling term d_0_ any peptide or protein with length ≤21 is subject to identical scaling while any peptide or protein >21 amino acids is scaled according to its length. For most of the peptides in this dataset this would make the score very sensitive towards small deviations and penalizes structural deviation of these peptides significantly more than longer ones. Furthermore, the lDDT, by definition, measures local deviations by considering only contacts within a radius of 15 Å. While this is favorable for measuring peptide positions that enter the binding pocket, it is less applicable for larger protein ligands that form tertiary structures distant to the receptor binding interface. In consequence, this could lead to a scenario where the prediction of a protein ligand retains most of its tertiary structure, i.e. moderate to high lDDT, but fails to predict the correct global placement, i.e. a high RMSD. Lastly, we find that the DockQ score is most suitable for the task at hand as it takes into account ligand RMSD (LRMSD), interface RMSD (iRMSD) and the fraction of retained native contacts (fnat), covering both local deviations with respect to the receptor (iRMSD and fnat), as well as global orientation (LRMSD).

### AlphaFold2 Initial Guess

AF2IG was used as described in https://github.com/nrbennet/dl_binder_design. Receptor-peptide dimers were preprocessed so that the peptide chain was the first chain of the file to ensure that AF2IG identifies ‘target’ chain and ‘binder’ chain correctly. The AF2IG script was modified to save the entire PAE matrix and to accept custom seeds from command-line. Seeds 1-50 were used for the predictions with `model_2_ptm.npz` weights (weights `model_1_ptm` to `model_5_ptm` were tested initially with `model_2_ptm` showing highest recovery; data not shown).

### Boltz-2

Boltz-2 was used with default settings apart from the `--use_potentials` flag. Initial tests considered both options for this flag but no significant difference could be observed (data not shown). No MSAs were allowed for either of the chains (i.e. `msa: empty`). For the receptor chain the respective reference structure was provided (‘apo’, i.e. peptide chain removed). Contrary to AF2IG and RF3, where the receptor chain residues were taken from the template, including gaps in the crystal structure, Boltz-2 requires a sequence. We opted to provide the full-length sequence taken from uniprot.org (58). Seeds 1-50 were used for the predictions with weights downloaded 06/20/25.

### RosettaFold3

To prevent RF3 from aborting low-confident runs `early_stopping_plddt_threshold` was set to 0.1. Other settings remained default. Analogous to Boltz-2, the apo version of the reference dimer was provided as prediction component. Seeds 1-50 were used for predictions with weights `rf3_latest.pt` obtained 08/15/25.

### BindCraft

Out of the three generative methods tested, BindCraft represented an outlier in that it follows an optimization protocol that aborts trajectories based on a variety of filters while BoltzGen and RFdiffusion3 rely on separate post-analysis to filter out unwanted designs. In order to ensure comparability with respect to the actual space sampled we set `Average_Binder_RMSD` and all stage-specific `Binder_RMSD` filters to null. Furthermore, we set `Average_pLDDT` and `1_pLDDT` and `2_pLDDT` filters to 0.5. Remaining filters were taken from the `peptide_filters.json` and kept unchanged. Advanced settings were taken from the `peptide_3stage_multimer.json` with `predict_initial_guess` and `enable_mpnn` set to `false`. Lastly, we collected both from Accepted as well as Rejected trajectories until the desired number of 10000 designs was met.

### BoltzGen

BoltzGen was used with the `peptide-anything` protocol while omitting all steps but the `design` step. Input structures were renumbered and hotspot residues adapted accordingly. Weights were cached 12/04/25.

### RFdiffusion3

RFdiffusion3 was used with `infer_ori_strategy: hotspots` where the `TIP` atoms were used for each of the defined hotspots and `is_non_loopy=True`. Any other settings were kept default. Weights `rfd3_latest.ckpt` were obtained 12/03/25.

### ProteinMPNN

Sequence generation with ProteinMPNN was performed using `--model_type “protein_mpnn”` with weights `protein_mpnn_v48_030.pt` and `--temperature 0.05`. The entire peptide chain was allowed for design while the entire receptor chain was kept fixed. One single sequence was generated per backbone.

## Author contributions

Hannes Junker: Conceptualization, Data Curation, Formal Analysis, Investigation, Visualization, Writing – Original Draft Preparation, Writing – Review & Editing

Clara T. Schoeder: Conceptualization, Funding Acquisition, Project Administration, Supervision, Writing – Review & Editing

## Acknowledgement

We thank D. Rieger and J. Meiler for valuable discussion regarding conceptualization and analysis. We also thank M. Beining for assistance with various scripts used for the calculations and analysis in this work. We further want to thank M. Ertelt for the critical reading of the manuscript. Computations for this work were done (in part) using resources of the Leipzig University Computing Cluster. The authors gratefully acknowledge the computing time made available to them on the high-perfomance computer at the NHR Center of TU Dresden. This center is jointly supported by the Federal Ministry of Research, Technology and Space of Germany and the state governments participating in the NHR.

## Supplementary Information

**Supplementary Table S1:**
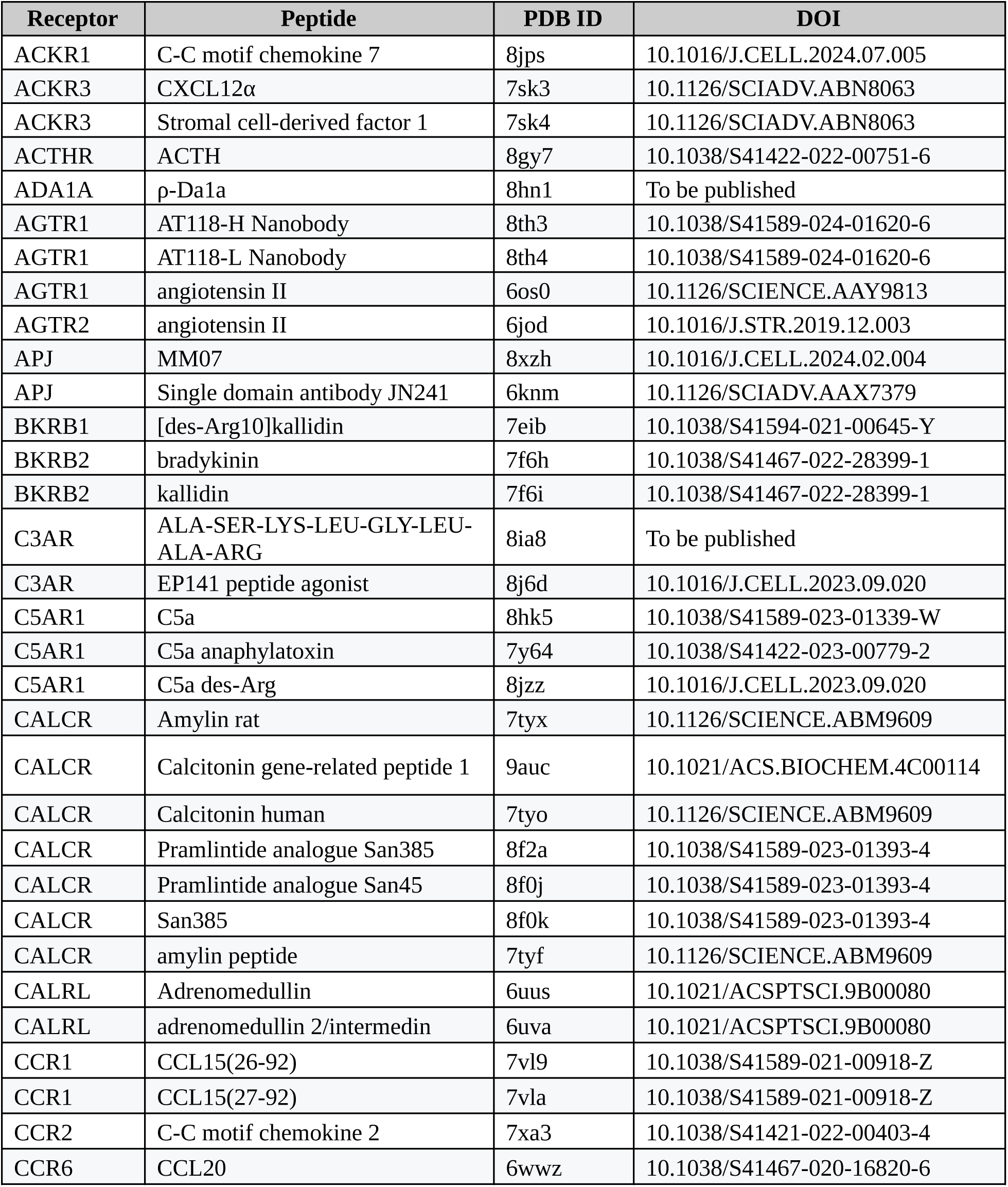
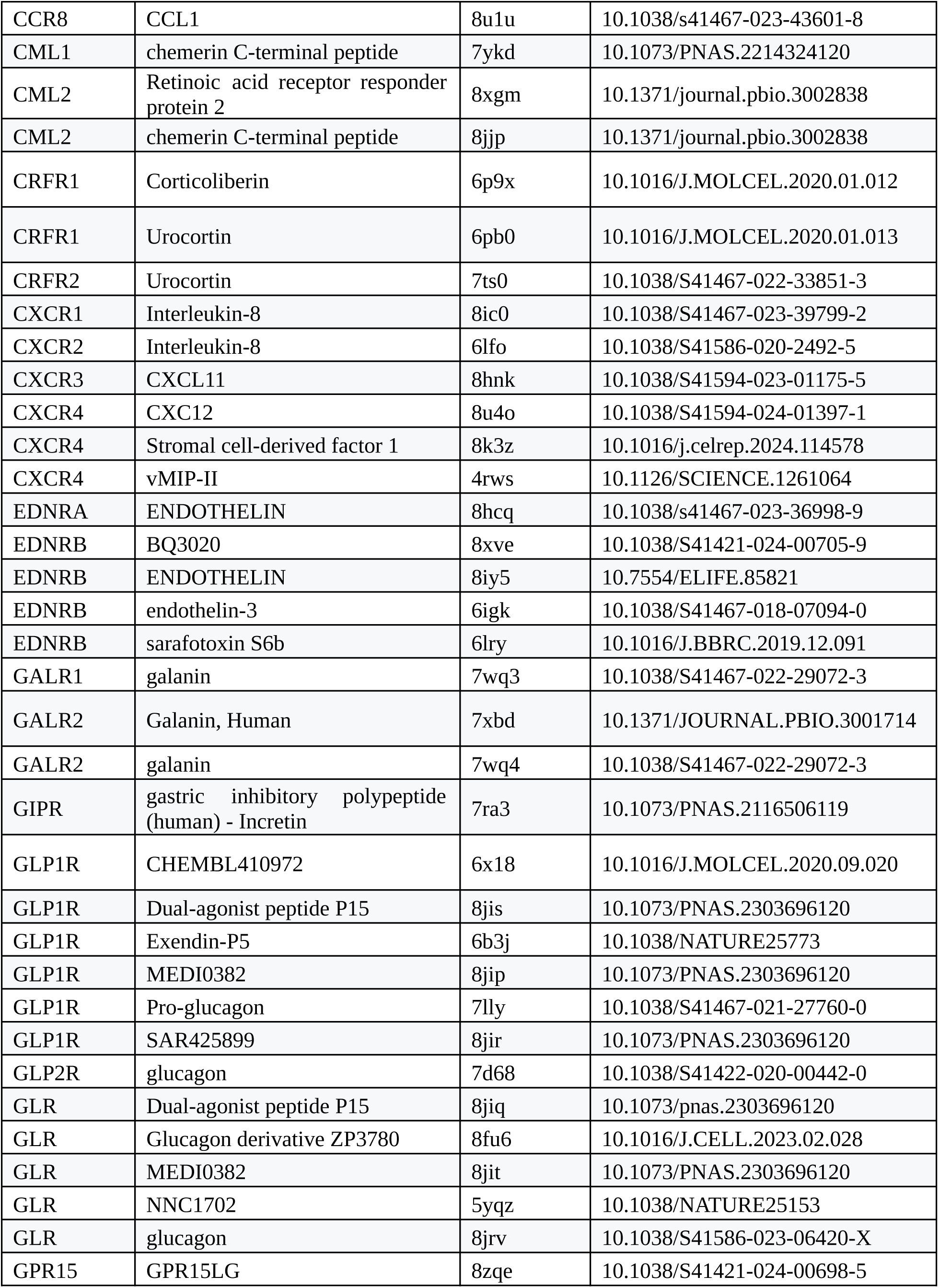
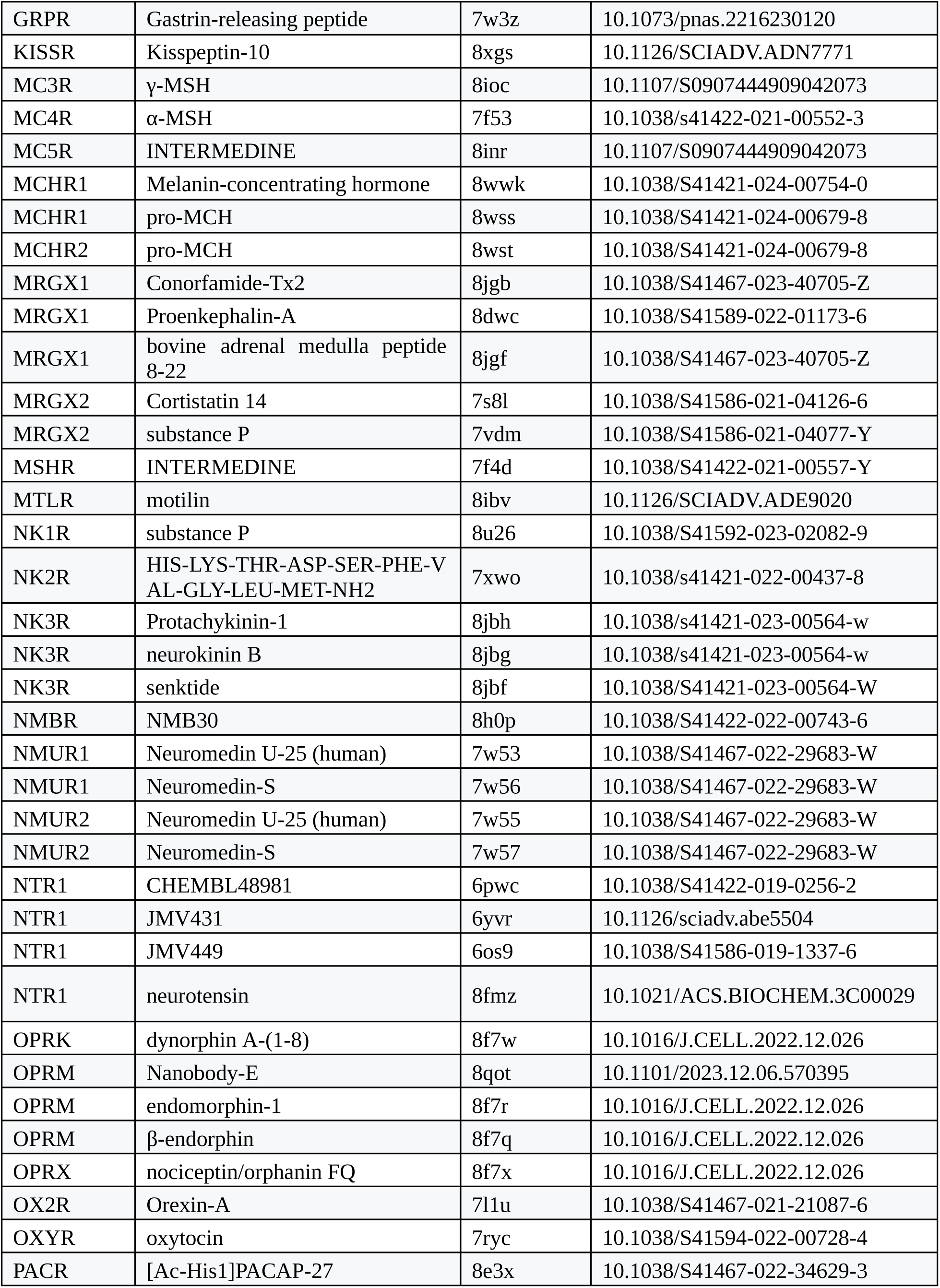
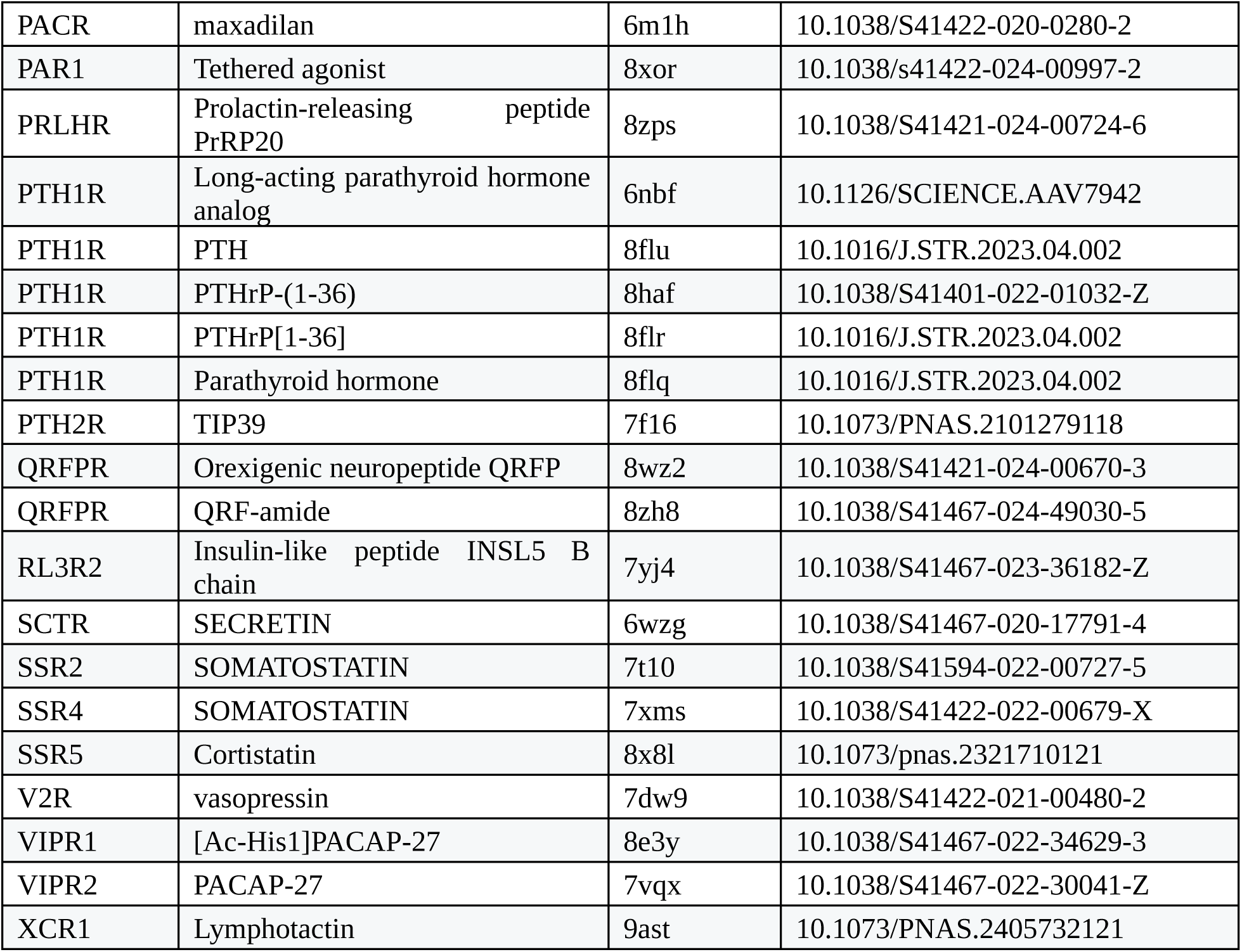
List of all 124 peptide-receptor complexes used to assess peptide prediction accuracy.

**Supplementary Fig. S1:**
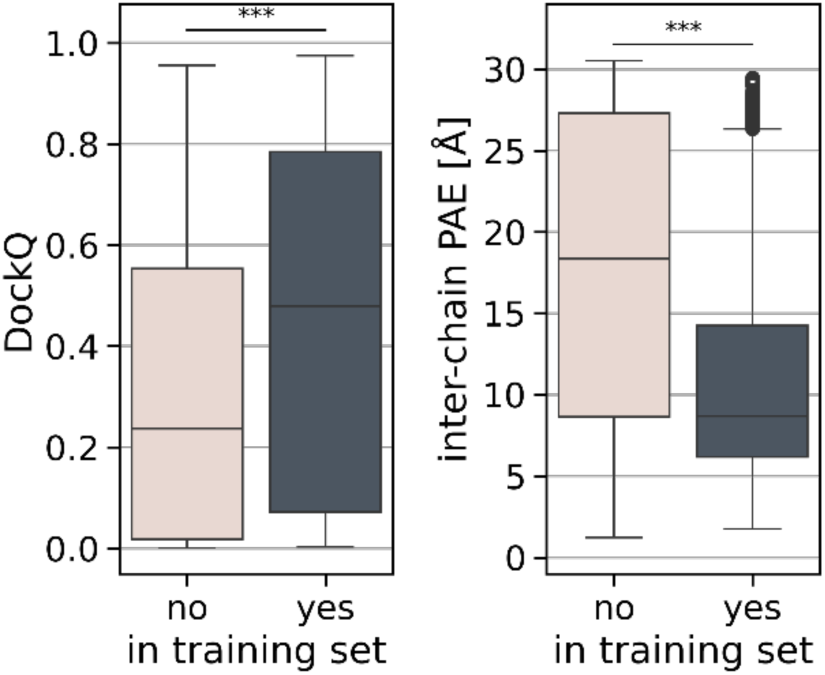
Both predicted structural similarity (left) and prediction confidence (right) decreases significantly for structures that were submitted after training date cutoffs. Statistical significance was assessed using Mann-Whitney U test (p<0.001 in both cases). A structure was deemed to be in the training set if the specific PDB ID or the same dimer with a different PDB ID was published prior to the respective training date cutoffs.

**Supplementary Fig. S2:**
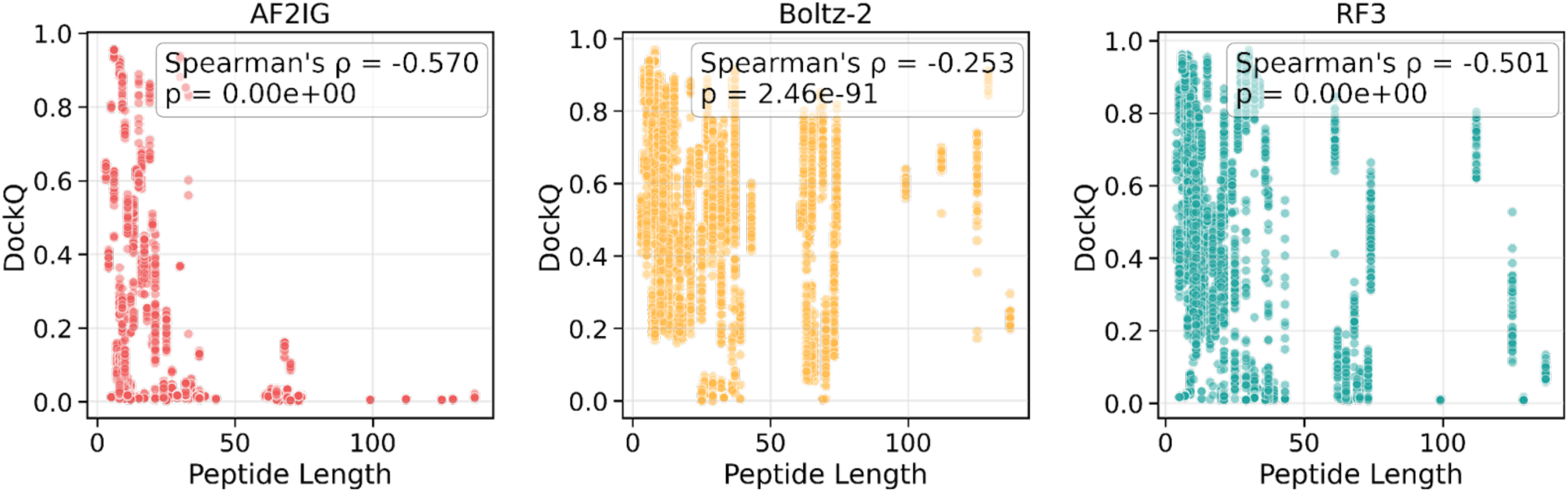
DockQ scores of every of the 50 predictions per receptor-peptide pair vs. peptide lengths.

**Supplementary Fig. S3:**
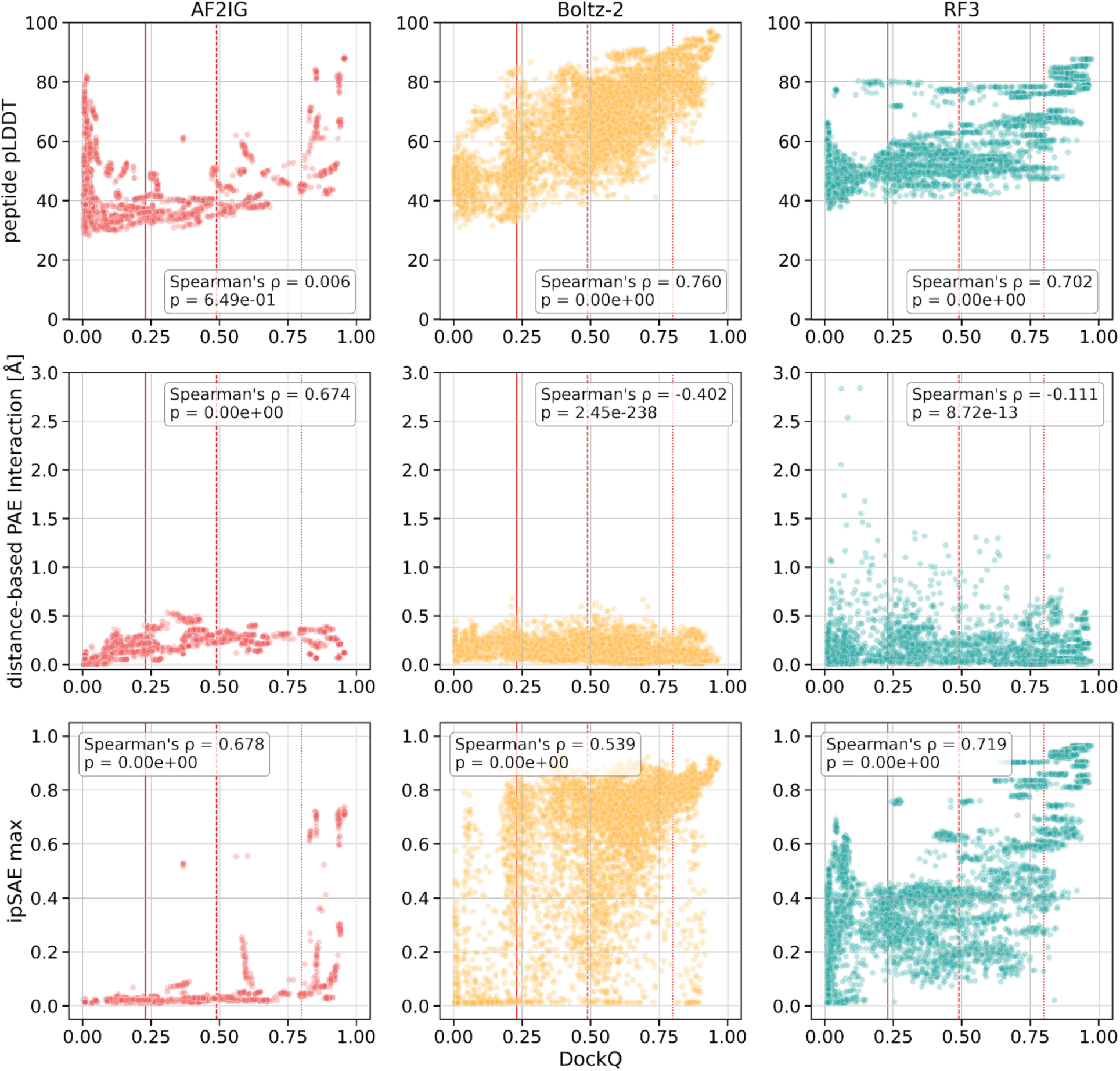
Additional confidence metric compared to DockQ scores by method for all predictions. Top row: Average pLDDT values for the peptide chain. Middle row: Distance-based PAE interaction; averaged PAE values for interchain residue pairs with a distance of ≤ 10 Å. Bottom row: iPSAE max calculated as in (44).

**Supplementary Fig. S4.**
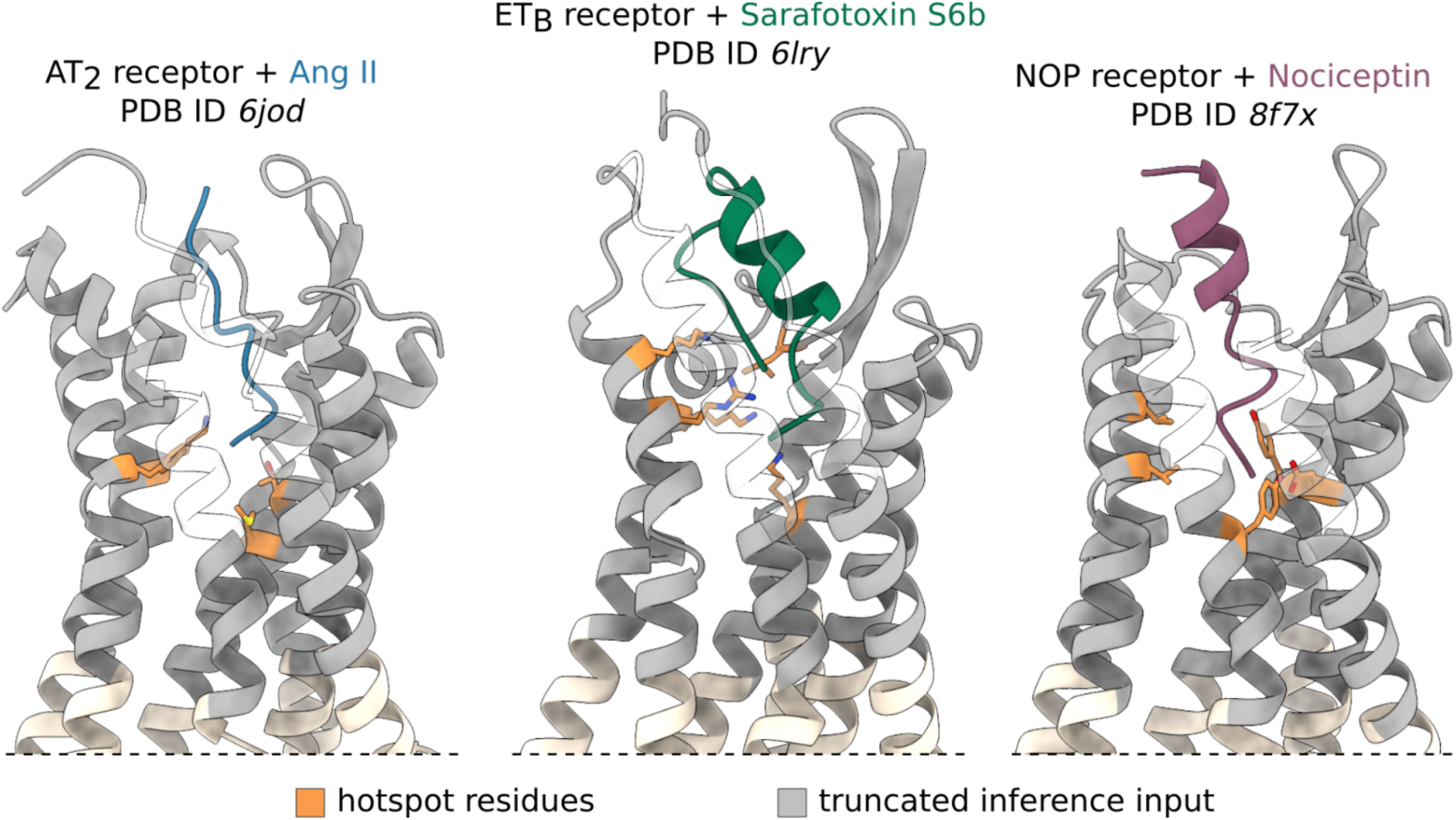
: Input structures for peptide generation. Hotspot residues highlighted in orange. Hotspots were selected based on minimum distance to the deepest pocket-penetrating residue. Starting from a minimum of three hotspots the final number of hotspots was determined in previous BindCraft test runs to ensure a reasonable amount of finished trajectories. Input structures (gray) were truncated to increase memory and runtime performance.

